# Trump Administration Impacts on Early Career Scientists and How To Fight Back

**DOI:** 10.64898/2025.12.03.691733

**Authors:** Crystal Hammond, John Patrick Flores, Siara Rouzer, Kassandra Fernandez, Amy Ralston, Adriana Bankston

## Abstract

Scientific research drives national progress and innovation, and must be protected at all costs. Actions from the Trump administration in early 2025 sent shockwaves across academic research. Federal science budgets were slashed, hundreds of research grants were abruptly frozen or terminated, and politically appointed agency heads overrode peer-reviewed grant decisions. We conducted rapid-response surveys to capture quotes, stories, and data to understand how these seismic shifts were being felt by early career researchers. The majority of survey respondents were PhD students, and early career trainees in varying years of training at a variety of institutions, mainly in the life sciences. Their future in research, funded by major science agencies including NIH, was threatened by funding cuts with varying impacts to their professional endeavors. Survey respondents highlighted concerns with long-term research job prospects and broader systemic issues in academia, leading them to consider leaving the field altogether and/or pursue science in another country. Their opinions highlighted concerns at the undergraduate, graduate, and postdoctoral levels. While universities provided some resources, trainees called upon Congress to protect, stabilize, and restore science funding. They also undertook a number of advocacy actions across the nation. The fight is far from over. Continued support for early career researchers and their careers in research is imperative to continue both in the current environment and into the future, in order to develop a strong workforce and ensure U.S. global competitiveness in science and technology fields.

## Current Landscape

Scientific research drives national progress and innovation, and must be protected at all costs. Defending scientific research requires fully funding it, as well as supporting the projects and careers of promising scientists who contribute to the STEM workforce and U.S. economic development. In early 2025, a surge of executive orders and the Trump administration’s FY 2026 budget proposal sent shockwaves across the academic research landscape, targeting everything from NIH extramural funding to climate science and DEI-related programs ^1,2^. Federal science budgets were slashed by double-digit percentages, hundreds of research grants were abruptly frozen or terminated, and politically appointed agency heads overrode peer-reviewed grant decisions ^3,4^. Not only will these actions lead to significant threats to research progress in the U.S., but it will impair our global competitiveness in the sciences.

As previously described, early March 2025 marked a turning point for researchers nationwide as these federal proposals cut science funding at agencies one by one ^5^. These measures severely impacted university research projects, talent recruitment and staffing in a number of ways. For example, on March 10, the University of Pennsylvania announced a campus-wide hiring freeze and a 5 percent cut in non-compensation expenses ^6^. By March 19, University of California (UC) System then-President Michael Drake ordered a system-wide freeze on all new faculty and staff searches and additional cost-saving measures. The UC also signaled a cautious approach to spending amid the funding uncertainty in response to the federal cuts ^7^. And on April 8, Northwestern University paused all hiring and cut discretionary costs by 10 percent ^8^. These are only a few examples highlighting the Trump administration’s negative impact on recruiting and retaining STEM talent in U.S. laboratories.

This dire situation has also impacted important grants supporting the next generation, including the NSF GRFP whose funding was cut in half in April 2025 (from about 2,000 to 1,000 in total) due the agency’s constrained budget ^9^. While this number was later partially restored by funding an additional 500 NSF GRFP grants ^10^, it now restricts second-year Ph.D. students from applying to a program that is critical to developing U.S. STEM talent ^11^. Additionally, funding to several National Institutes of Health (NIH) grants, including the IRACDA program, was cut by the Trump administration ^12^ and later restored by federal courts ^13^. And at major universities such as the UC System, current president James Milliken reversed the decision to end financial support for hiring postdoctoral fellows out of the UC President’s Postdoctoral Fellowship Program ^14^. Although there is restoration of some program funding to support the STEM pipeline, this unstable situation at major federal agencies is deterring significant early career talent from remaining in our research pipeline.

In May 2025, the Trump administration released its FY 2026 budget proposal calling for a 40 percent cut to NIH and a 55 percent cut to NSF ^15,16^, freezing new grants and stalling many projects overnight ^17^. This was documented by several groups including Association of American Universities (AAU) showcasing proposed research funding cuts from the administration across federal agencies ^18^. While only a few examples, these swift actions made it clear that a single federal proposal can stop vital research in its tracks and halt scientific progress at multiple agencies and research in institutions across the country, which also impacts student populations ^19^.

We do not yet have a full grasp of the long-term consequences of these actions on the U.S. research enterprise, but we can affirm that these actions will lead to a significant reduction of talent in the STEM workforce. As universities have begun to cut Ph.D. admissions ^20,21^, early career researchers are seeking roles elsewhere as job losses and grant terminations increase at U.S. institutions, meanwhile several other nations are heavily recruiting STEM talent into their regions resulting in talented scientists moving to other countries ^22–24^, particularly as they are investing heavily in innovations attracting the world’s brightest minds ^25^. In addition to impacts on the research pipeline, staff reductions and job losses experienced by scientific experts at our federal agencies, including at NSF, will heavily set back American leadership in science and technology by undercutting funding for federally-funded research, education and diversity initiatives ^26^. Cutting indirect costs at the NIH also has severe consequences on American science and innovation ^27^. Without dedicated federal research funding, or dedicated agency staff to manage these grants, early career researchers won’t be able to get their ideas off the ground and build fruitful science careers across U.S. laboratories.

## Surveys Conducted

To understand how these seismic shifts are being felt on the ground, especially by early career researchers at a critical juncture in their scientific paths, we conducted a rapid-response survey to capture their quotes and stories, including any advocacy actions taken (survey #1, **appendix**), with an additional data-driven survey to learn about impacts of federal funding cuts on their research and careers, and on the broader academic research workforce (survey #2, **appendix**). The surveys were developed by members of the Sai Resident Collective (SRC) Bankston Group, a project of the Civic Science Media Lab ^28^ and in collaboration with the non-profit Science for Good ^29^. Sai Resident Collective (SRC) is a non-profit incorporated in Massachusetts as the Stem Advocacy Institute.

The surveys were administered online using Google Forms between May 23 and June 6, 2025, in order to obtain a rapid-fire response from early career trainees in our networks while the Trump administration’s actions were unfolding. The first survey (survey #1) was brief and time-sensitive to capture these responses, with additional data-driven results from early career researchers across various disciplines and institutions across the country (survey #2). For both surveys, participants were recruited through academic listservs, social media, and direct outreach by SRC residents and collaborators. Survey participation was voluntary and opt-in anonymous for personally identifiable information, with initial results from survey #1 previously documented. Survey respondent names were only included in the publication and prior blog post ^5^ with explicit consent, otherwise anonymous.

## Results and Brief Discussion

In this publication, we show results from the data-driven survey (survey #2) (30 responses), including additional views from the quotes and stories survey, including any advocacy actions taken by respondents (survey #1) (7 responses including Atlantic International University) to highlight the landscape of our findings. The data-driven survey yielded early career responses to questions related to career stage, time in current position, institution, field of study, demographics information (optional), federal agency funding current research, impacts of funding cuts on research careers including specific details, concerns related to research projects and career, considering leaving academic research, resources provided by the university, and what Congress could do to support.

## Career Stage and Time in Current Position

On the career stage of respondents (Q1), the majority were PhD students (60%), followed by undergraduate students (13.3%), postdoctoral researchers (13.3%) and an “other” category (13.3%), with no Master’s student responses (**Figure 1**). At the time when the survey was conducted, respondents had been in their current positions (Q2) for 1-3 years (43.3%), followed by 4-5 years (40%) and 5+ years (16.7%) (**Figure 2**). The majority of responses coming from PhD students and trainees from 1-5 years in their positions is significant given the contributions of this population to the workforce, and their uncertain scientific futures in the current environment.

**Figure 1.**
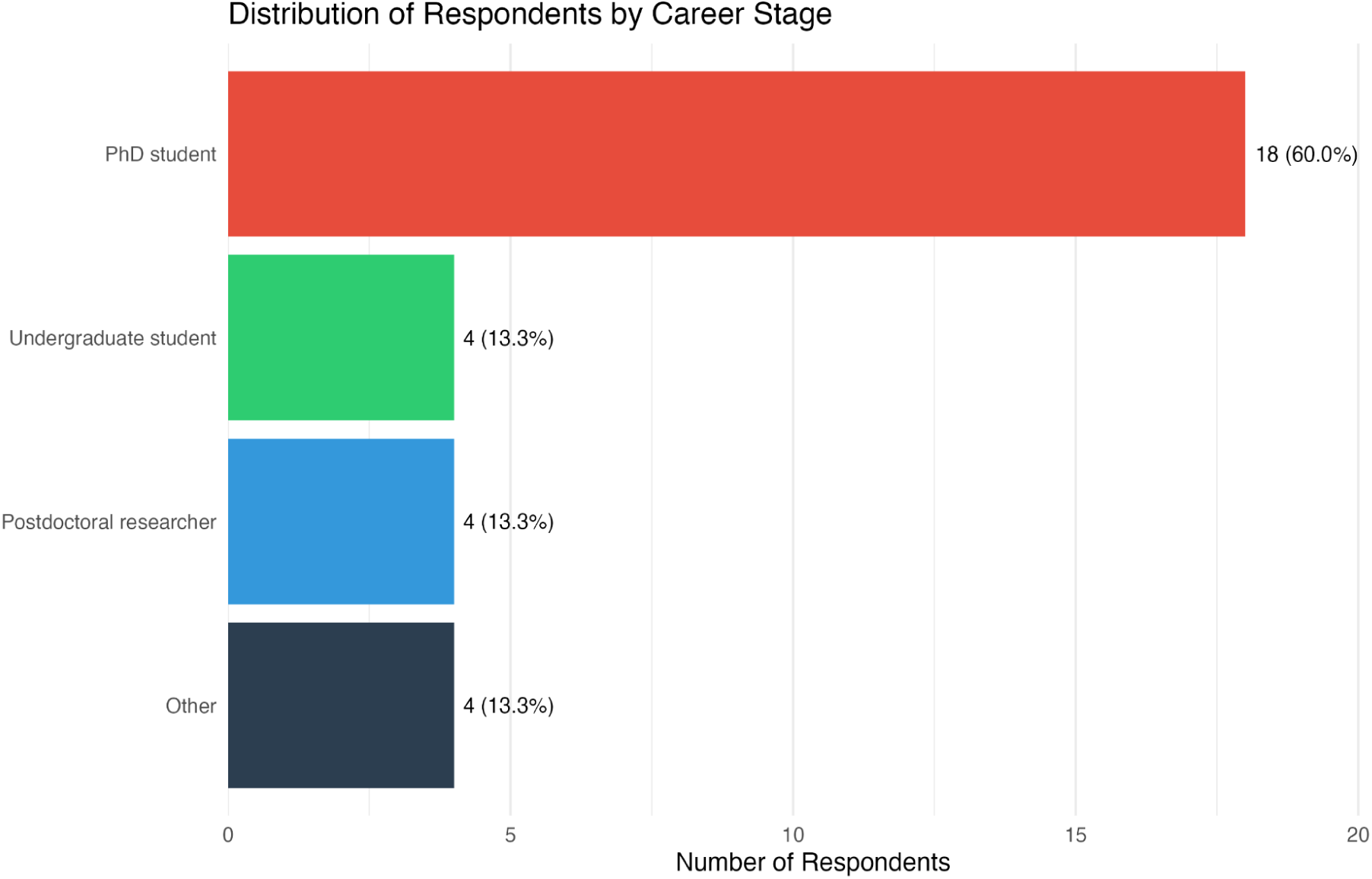
Distribution of Respondents Career Stage (Q1). The majority of respondents from survey #2 were PhD students (60%), followed by undergraduate students (13.3%), postdoctoral researchers (13.3%) and an “other” category (13.3%), with no Master’s student responses.

**Figure 2.**
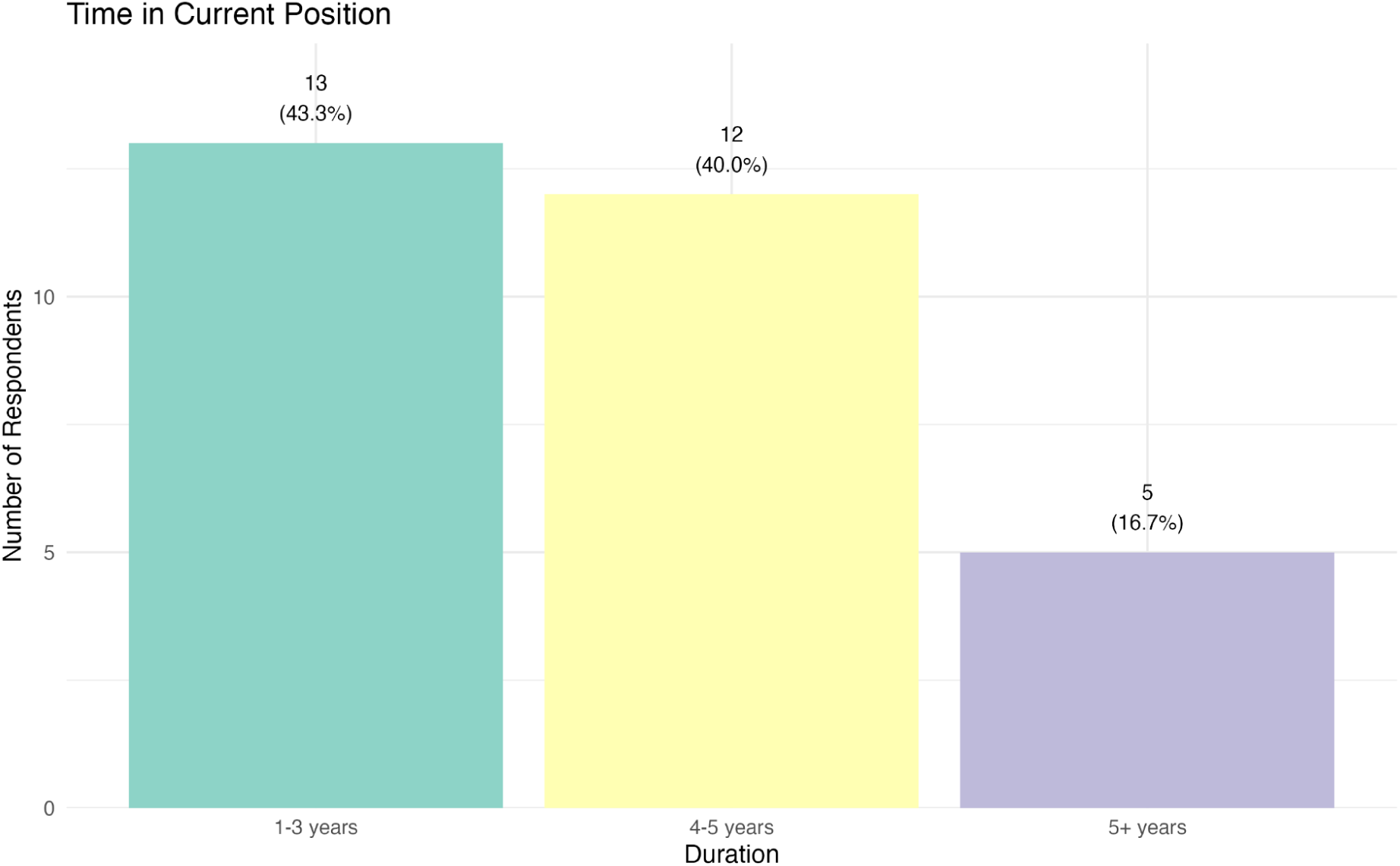
Respondents Time in Current Position (Q2). Respondents from survey #2 were in various career stages, and in their current positions for 1-3 years (43.3%), followed by 4-5 years (40%) and 5+ years (16.7%).

## Academic Institution and Field of Study

Respondents wrote in the academic institution where they were studying or working at the time of the survey (Q3), which resulted in the table below. We then classified these data into public and private schools with additional details using the Carnegie Classification information. These were R1 - Very High Spending and Doctorate Production (∼spending $50 million on research & development and awarding at least 70 research doctorates), R2 - High Spending and Doctorate Production (∼spending at least $5 million on research & development and award at least 20 research doctorates), and finally Research Colleges and Universities (∼spending at least $2.5 million on research & development, not including R1 and R2 schools) ^30^ (**Table 1**, **Figure 3**). Upon analyzing information provided by academic institutions using independent criteria, we found that a large majority of respondents were from R1 schools (∼67%) and public institutions (∼67%), and that the largest share of institutions were located in the Northeastern U.S. (∼43%) (data not shown). Having a majority of respondents from R1 schools and public schools could be a result of outreach, and/or a desire for those respondents to complete the survey in order to showcase impacts of the Trump administration on their research.

**Figure 3.**
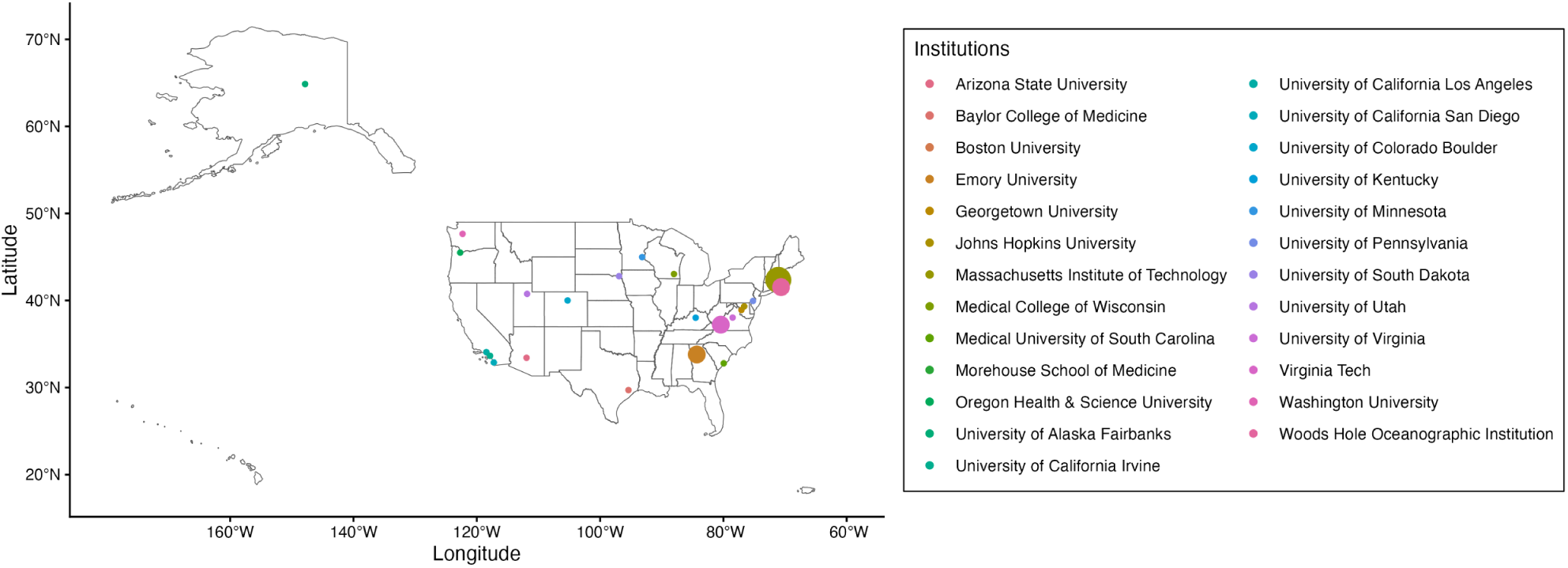
Map of Academic Institutions Represented by Survey Respondents (based on Q3). Data from this map is from both conducted surveys, with self-reported respondent institution data from table 1. The larger dots denote a greater number of responses. Atlantic International University was not included.

**Table 1.** Academic Institutions Represented by Survey Respondents (Q3). Academic institutions reported by respondents, which we then classified as public and private schools with additional details using the Carnegie Classification resulting in R1, R2, or Research Colleges and Universities. This table is from both conducted surveys with self-reported respondent data on institutions, including the data-driven survey (entire list) (survey #2) and additional quotes and stories survey (survey #1) (**). Atlantic International University was included in the survey responses which we retained in this table for completion of the responses provided. We later discovered that this is not an accredited university ^31^ and therefore did not include it in Figure 2.

| University Represented by Respondents | Public/Private University | Carnegie Classification |
| --- | --- | --- |
| Atlantic International University** | Private | N/A |
| Arizona State University** | Public | R1 |
| Baylor College of Medicine | Private | R1 |
| Boston University | Private | R1 |
| Emory University** | Private | R1 |
| Georgetown University** | Private | R1 |
| Johns Hopkins University | Private | R1 |
| Massachusetts Institute of Technology | Private | R1 |
| Massachusetts Institute of Technology<br>+ Woods Hole Oceanographic Institution | Private | R1 + N/A |
| Medical College of Wisconsin | Private | R2 |
| Medical University of South Carolina | Public | R1 |
| Morehouse School of Medicine (recent<br>graduate)** | Private | Research College or University |
| Oregon Health & Science University | Public | R2 |
| University of Alaska Fairbanks** | Public | R2 |
| University of California Irvine | Public | R1 |
| University of California Los Angeles | Public | R1 |
| University of California San Diego | Public | R1 |
| University of Colorado Boulder | Public | R1 |
| University of Kentucky | Public | R1 |
| University of Minnesota - Twin Cities<br>+ MN Sea Grant | Public | R1 + N/A |
| University of Pennsylvania | Private | R1 |
| University of South Dakota | Public | R2 |
| University of Utah | Public | R1 |
| University of Virginia | Public | R1 |
| Virginia Tech | Public | R1 |
| Washington University in St. Louis | Private | R1 |

A map was generated in R by geocoding the institutions which survey respondents provided (in Table 1, with the exception of Atlantic International University) and overlaying this information as colored dots on a U.S. map, with larger dots denoting a greater number of responses (**Figure 3**). Our results showcase important impacts on early career researchers, hindering their ability to contribute to the U.S. workforce and competitiveness due to reduced federal science funding for their projects and careers in science and technology fields in this country.

In terms of disciplines (Q4), the majority of respondents were in the life sciences (63.3%), followed by engineering (16.7%), other (10%), environmental sciences (6.7%) and social sciences (3.3%) (**Figure 4**). This is unsurprising given the generally high prevalence of early career scientists performing laboratory research in life sciences, thus a relatively large segment of the U.S. research workforce, many of whom are considering leaving the country ^32^.

**Figure 4.**
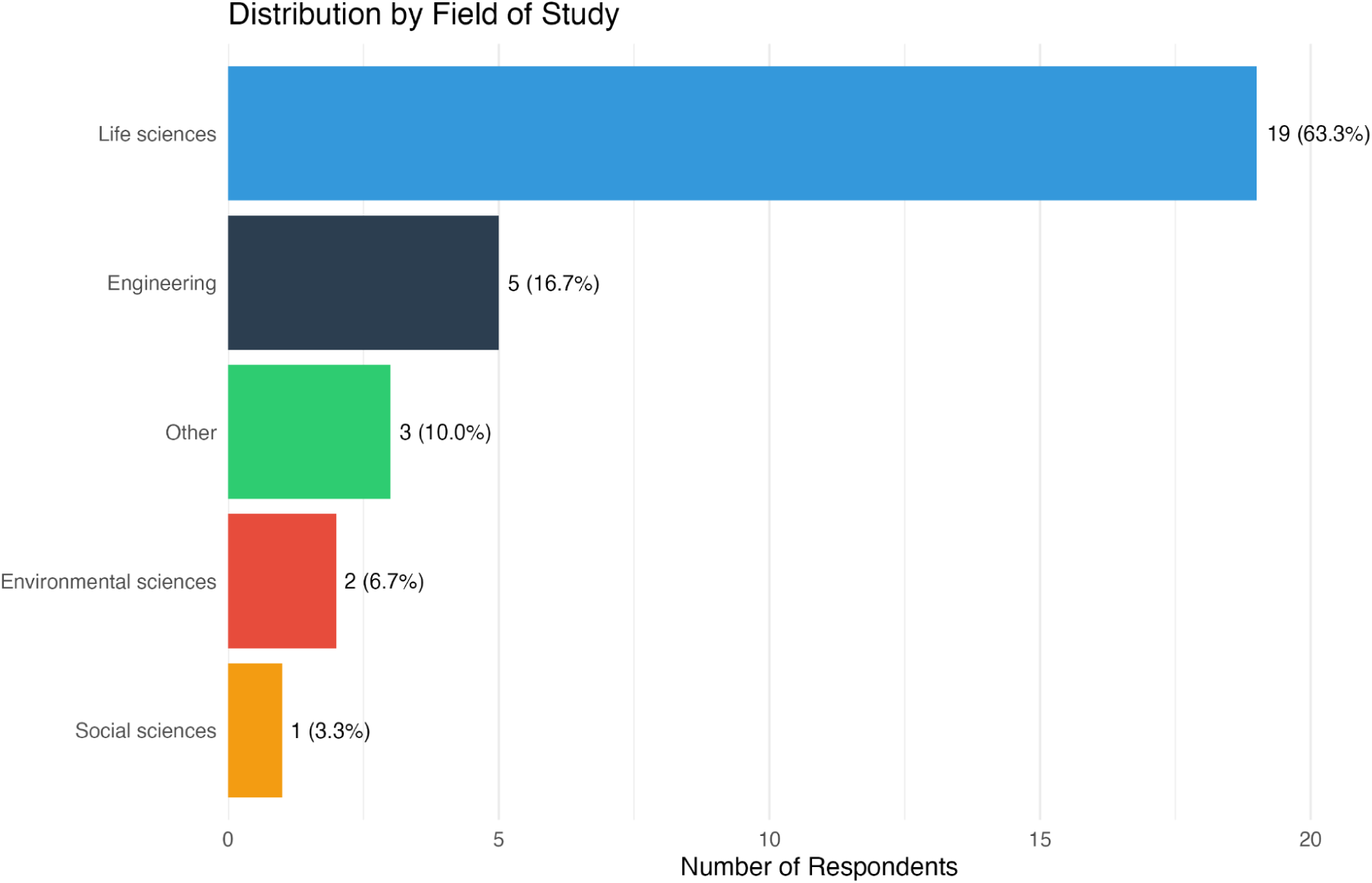
Respondents Academic Institution and Field of Study (Q4). Respondents from survey #2 were in various fields of study, with a majority in the life sciences (63.3%), followed by engineering (16.7%), other (10%), environmental sciences (6.7%) and social sciences (3.3%).

## Racial Identity Data

Although the demographics question was optional (Q5), all the participants opted to respond, with responses consisting in White (70%), followed by Asian (13.3%), Black or African American (6.7%), Bi or Multi-racial (6.7%), and American Indian or Alaska Native (3.3%), with no responses in the Native Hawaiian or Other Pacific Islander and “Other” categories (**Figure 5**). The highest rate of responses from White individuals may be reflective of their comfort in completing such a survey as compared to other groups targeted by the Trump administration.

**Figure 5.**
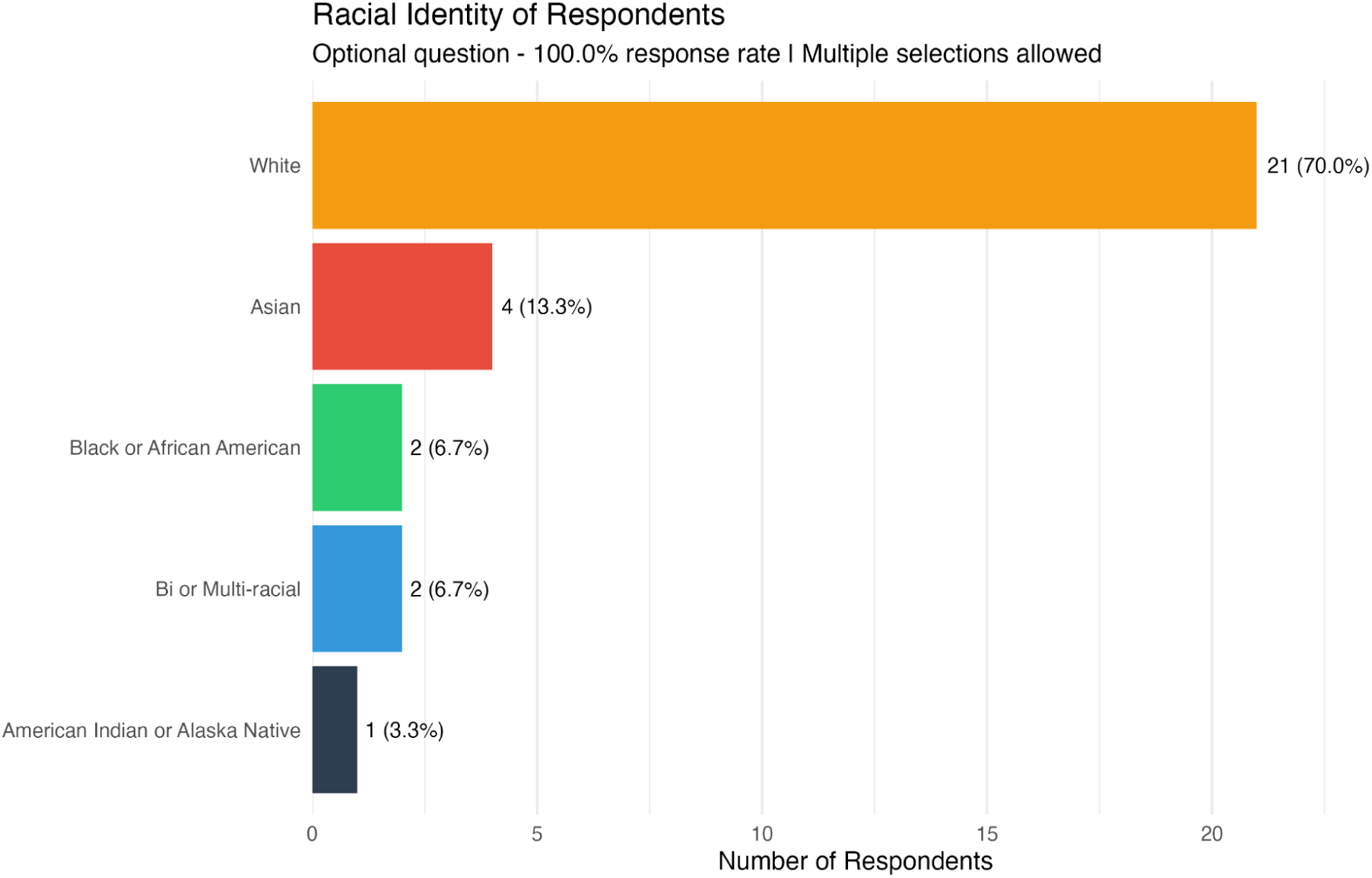
Racial Identity of Respondents (Q5). Respondents from survey #2 were majority White (70%), followed by Asian (13.3%), Black or African American (6.7%), Bi or Multi-racial (6.7%), and American Indian or Alaska Native (3.3%), with no responses in the Native Hawaiian or Other Pacific Islander and “Other” categories.

## Federal Agency Funding for Research and Impacts of Recent Science Funding Cuts

The breakdown for the federal agency research performed by respondents (Q6) included as top choices NIH (63.3%), other (30%), NSF (20%), DOD (13.3%), USDA (10%), NIST (3.3%) and NASA (3.3%), with no DOE option selected (**Figure 6**). Broadly, these data indicate the importance of federal funding at these agencies for supporting research breakthroughs.

**Figure 6.**
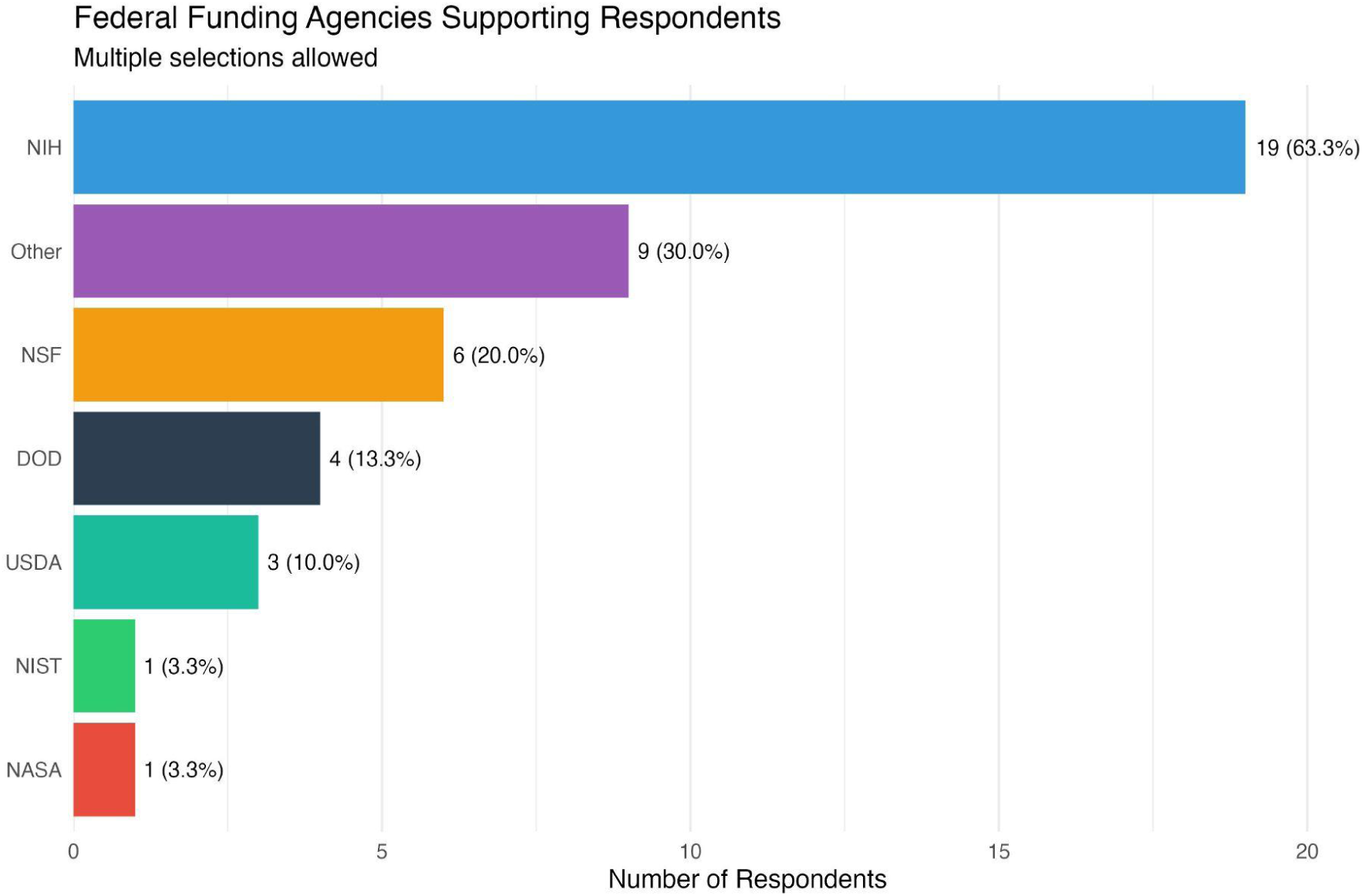
Federal Funding Agencies Supporting Respondent Research (Q6). Respondents from survey #2 indicated NIH as the top research funder (63.3%), followed by other (30%), NSF (20%), DOD (13.3%), USDA (10%), NIST (3.3%), NASA (3.3%), with no DOE option selected.

When given a scale of 1 - 10 (1 = No effect, 5 = Moderately affected, 10 = Severely affected) to rank how much the recent funding science cuts have affected them professionally (Q7), respondents have varied responses we then batched into low impact (scores 1-3), moderate impact (scores 4-7) and high impact (scores 8-10) for easier visualization. With this visualization, respondents indicated 10% low-impact, 40% moderate impact, and 50% high impact of federal funding cuts on their professional endeavors (**Figure 7**). Given the majority responses in the moderate or high impact severity categories, these data indicate detrimental effects resulting from these cuts on the U.S. research talent and workforce.

**Figure 7.**
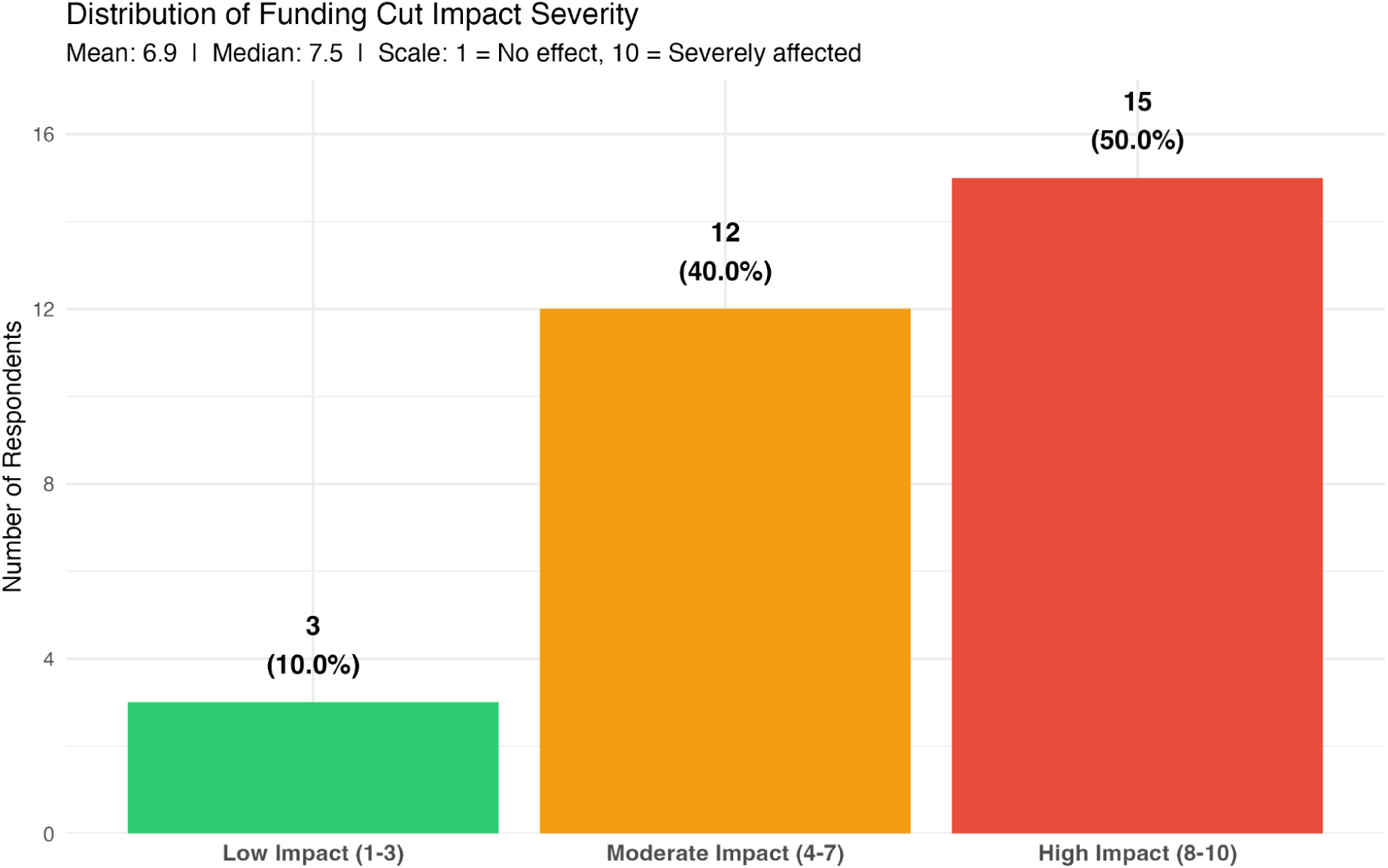
Distribution of Impact Severity of Federal Research Funding Cuts (Q7). On a 1-10 scale (1 = No effect, 10 = Severely affected), self-reported answers highlighted 10% of respondents indicating low impact severity (scores 1-3), 40% indicating moderate impact severity (scores 4–7), and 50% indicating high impact severity (scores 8–10).

## Professional Impacts of Science Funding Cuts, Biggest Concerns, Concerns with Impacts on Research and Careers & Leaving Academia

When asked about impacts experienced based on university actions, respondents (Q8) indicated besides “other” responses (53.3%), that their stipend and benefits were cut or delayed (20%), for some nothing had changed (16.7%), and for others the university froze hiring and/or rescinded the offer (10%) (**Figure 8**). While we may be missing some important data given the high percentage of respondents in the “other” category, these results overall indicate detrimental professional impacts on careers of early career scientists. In a broader sense, respondents highlighted concerns with potential future funding cuts on their research and careers in science (Q9), included with long-term research job prospects (90%), with many considering pursuing research in another country besides the U.S. (60%), or concerns with an inability to complete their research project (56.7%), grants not being renewed (53.3%), no longer wanting to pursue a research career (46.7%), and other (26.7%) responses (**Figure 9**). These significant concerns will prevent early career researchers in being able to pursue science careers in the U.S. which is a threat to our competitiveness on the global stage.

**Figure 8.**
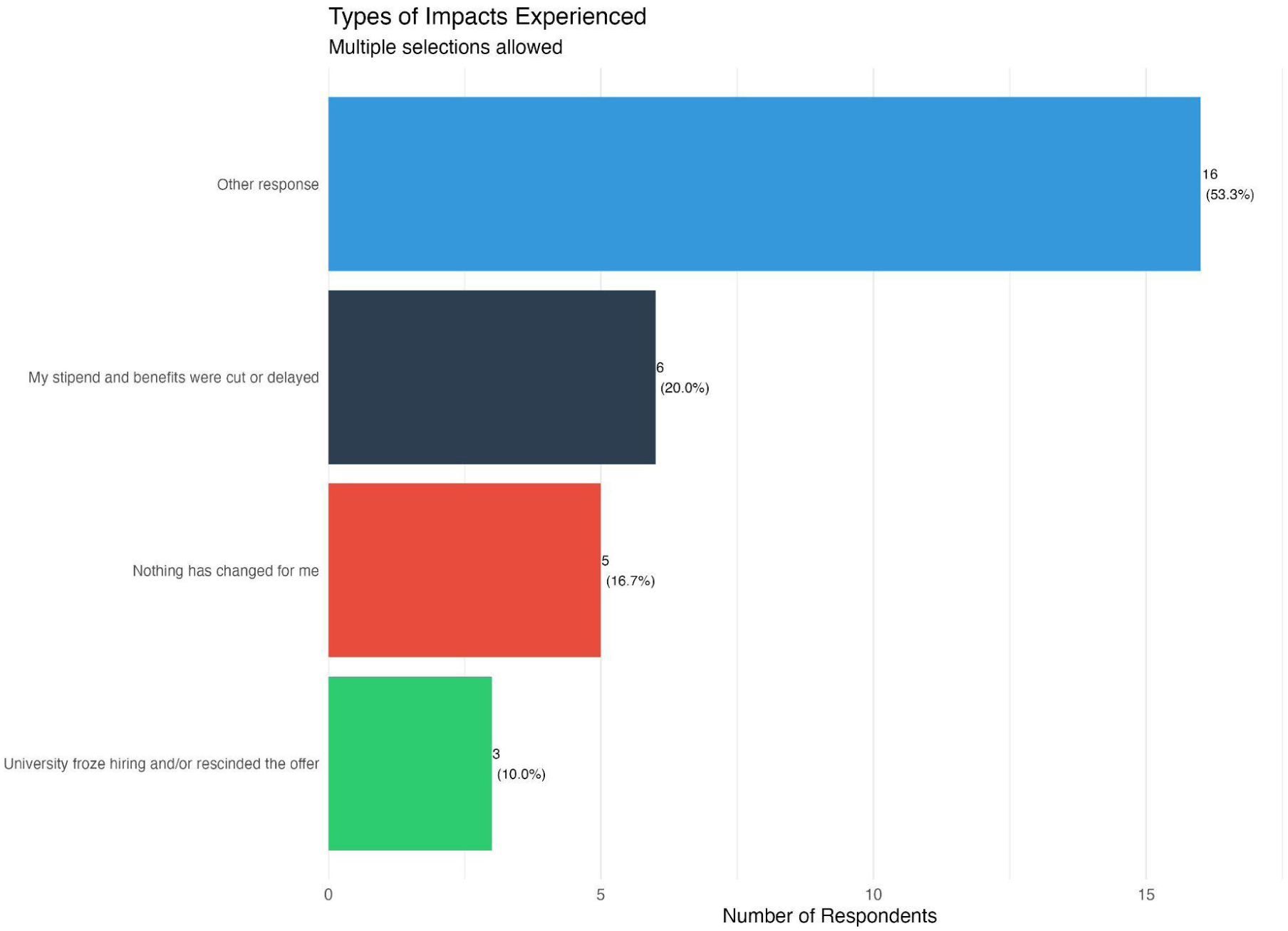
Professional Impacts Experienced Resulting from Science Funding Cuts (Q8). Respondents from survey #2 indicated besides “other” responses (53.3%), that their stipend and benefits were cut or delayed (20%), for some nothing had changed (16.7%), and for others the university froze hiring and/or rescinded the offer (10%).

**Figure 9.**
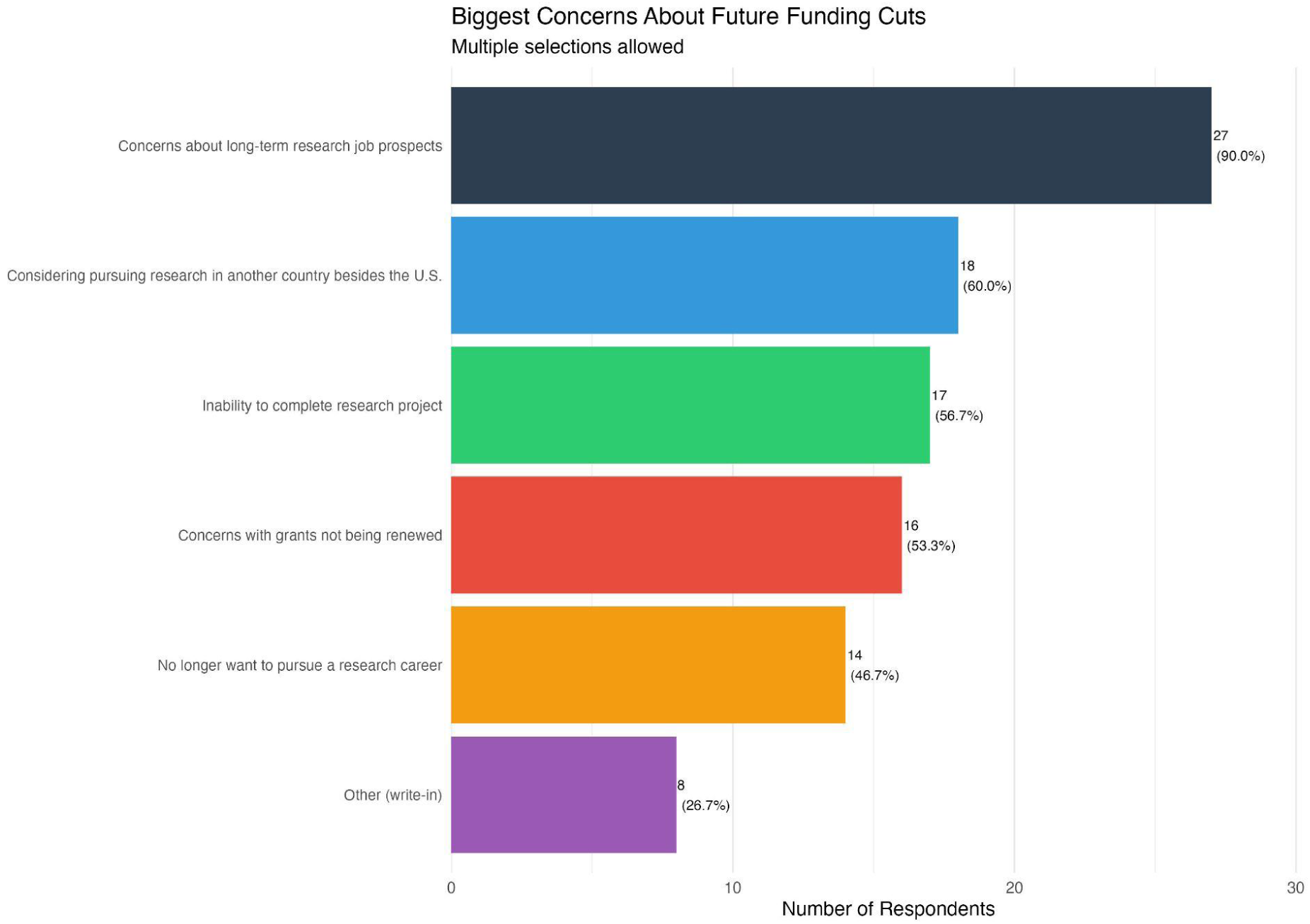
Biggest Concerns About Future in Science Due to Science Funding Cuts (Q9). When asked about impacts on research and careers in science, respondents from survey #2 indicated concerns about long-term research job prospects (90%), considering pursuing research in another country besides the U.S. (60%), inability to complete research project (56.7%), concerns with grants not being renewed (53.3%), no longer want to pursue a research career (46.7%), and other (26.7%).

At the systemic level, respondents highlighted particular areas of professional impact due to federal funding cuts (Q10), with the highest effects being on mental health and wellness (73.3%), followed by education and training (63.3%), belonging in STEM (56.7%), pay and benefits (40%), and other (6.7%) (**Figure 10**). When asked if they were considering leaving academic research, many said yes (56.7%), some said no (26.7%), with other (16.7%) responses (**Figure 11**). These data show negative impacts on stipends, benefits, hiring and offers for early career scientists, with long-term consequences on research projects and careers, including grant renewals. Moreover, these responses showcase detrimental impacts beyond research projects, extending to aspects critical for early career trainees to thrive in their careers including several systemic issues leading to loss of U.S. talent.

**Figure 10.**
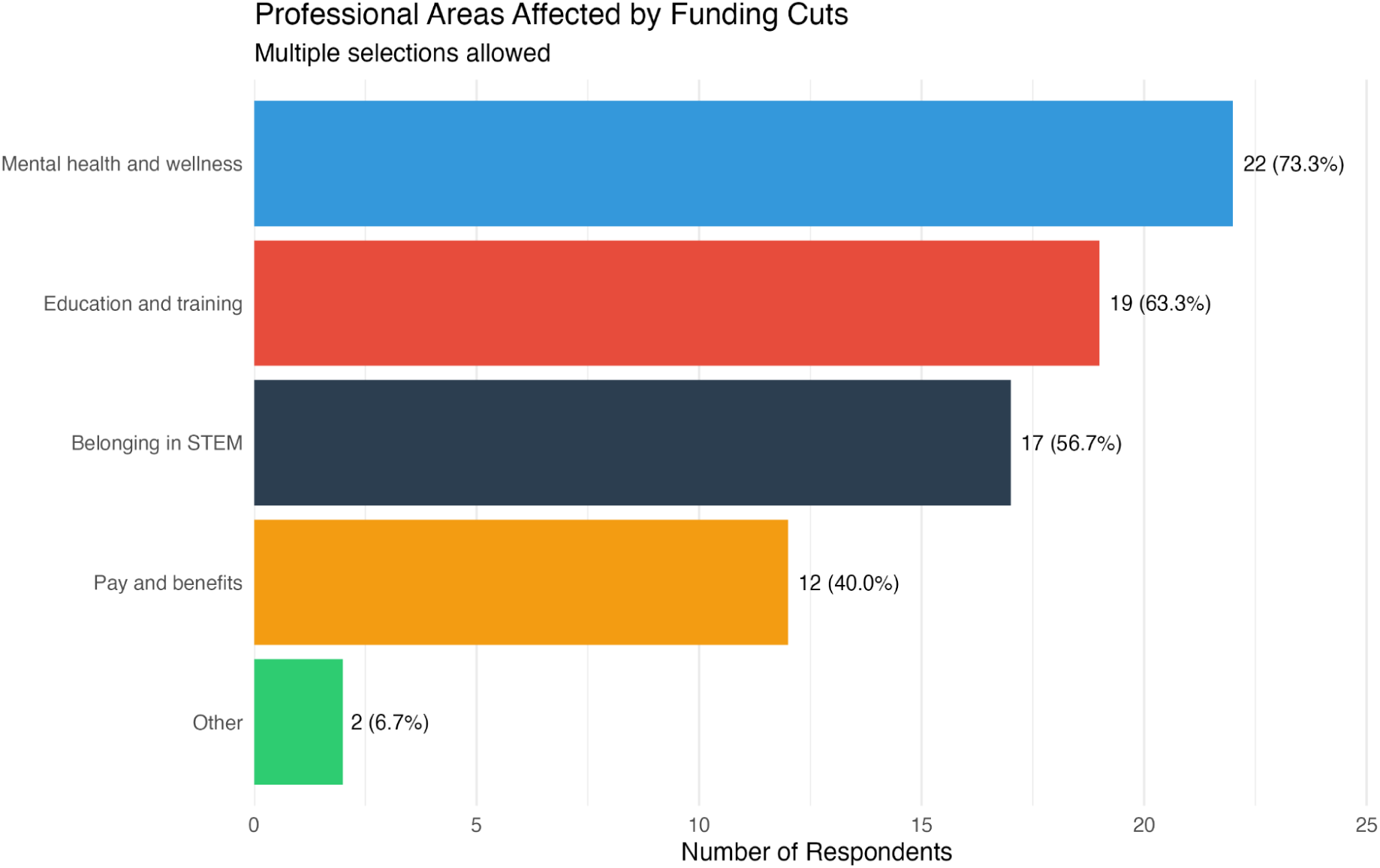
Impacts of Federal Funding Cuts on Respondents Research and Careers (Q10). Respondents from survey #2 indicated systemic issues resulting from federal funding cuts, including mental health and wellness (73.3%), education and training (63.3%), belonging in STEM (56.7%), pay and benefits (40%), and other (6.7%).

**Figure 11.**
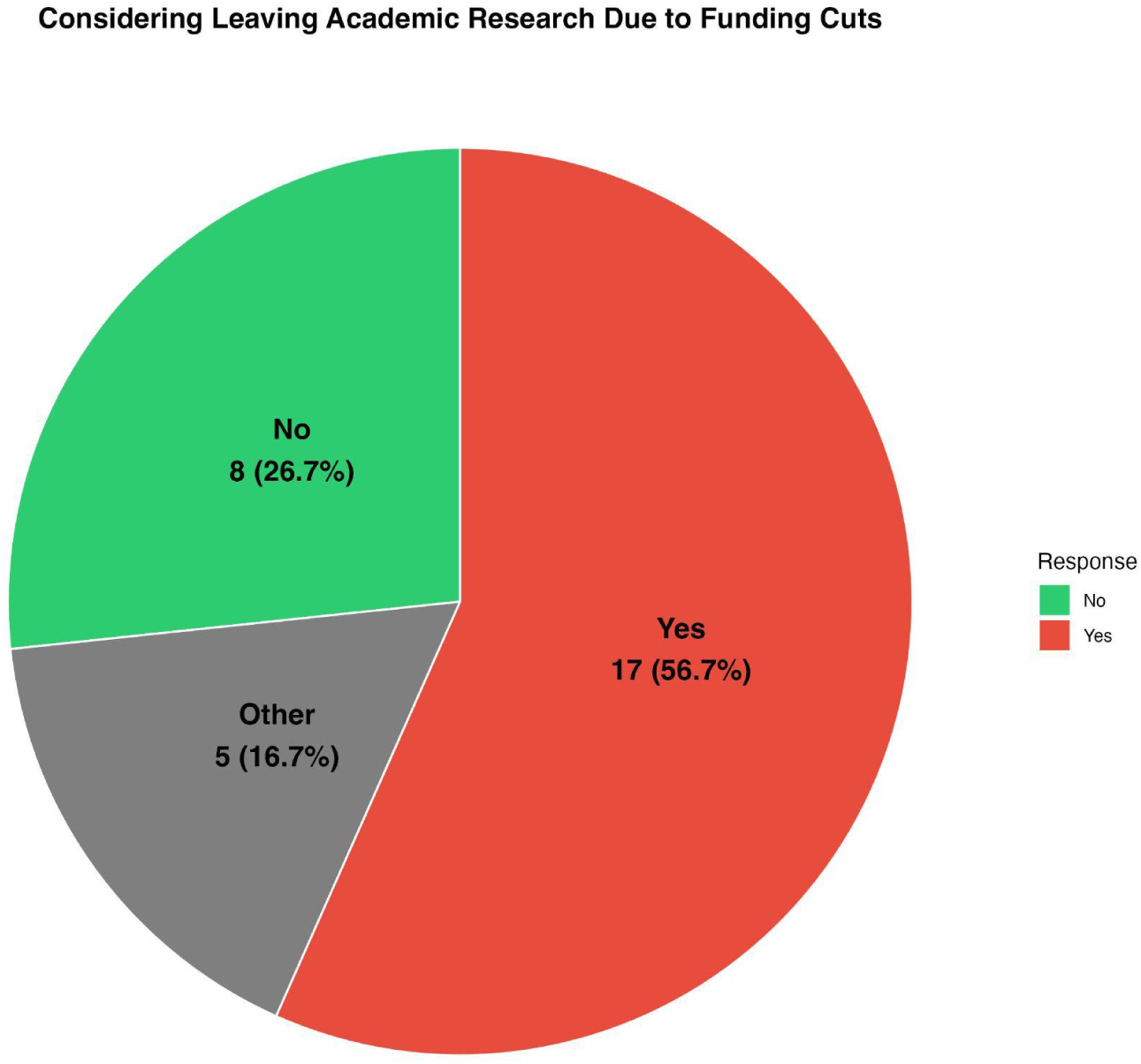
Respondents Considering Leaving Academia Due to Science Funding Cuts (Q11). Respondents from survey #2 indicated the majority (56.7%) are considering leaving academic research, whereas some were not (26.7%) with other (16.7%) responses.

## Quotes from Survey Respondents (Surveys #1 and #2)

In addition to data-driven results, we also collected a number of quotes from trainees to elevate their voices and enable them to share thoughts on the Trump administration’s actions on their careers and longer-term future in science, including universities and national laboratories. These opinions are gathered from both surveys (surveys #1 and #2), with a longer description showcased from those who wanted to share more detailed views.

### Undergraduate Students: Cutting Off the Source

While we primarily intended to capture views from graduate students and postdoctoral researchers, undergraduate students also responded and we wanted to document their views. This is important as many of them might seek to attend graduate school and pursue research careers long-term. With funding sources cut-off for undergraduate research, they may choose to pursue other avenues, which could include pursuing college and/or education in other countries ^33^. This is likely the case due to funding cuts to the NSF REU program, which many talented students utilize to explore and begin their research careers ^34^. An **anonymous** undergraduate student from University of Alaska Fairbanks who is pursuing a B.S. in wildlife biology in their 4^th^ year at the time of the survey, states that her “first undergraduate research project was canceled by [her] university because of federal funding issues despite [her] research being fully funded by the department of veterans.” As a disabled veteran, these include broader impacts on her mental health, leading her to affirm that she is “physically disabled” and calling for the ending of “disability discrimination” in STEM fields within universities. While this is only one example and a larger systemic issue, this opinion is unlikely to be unique among undergraduate students who may be turned away from staying in science due to the Trump administration’s actions and related university policies that hinder their ability to thrive in their research and education.

### Graduate Students: Dreams on Hold, Futures in Flux

Across disciplines, graduate trainees ^35^ describe a sudden dislocation, where their carefully-timed dissertations, job searches, and sense of scientific purpose have been knocked off-course by the 2025 federal funding cuts shock. We surveyed these students and showcased their responses. **Spencer Waters**, a neuroscience PhD candidate at Georgetown University, reports that “illegal funding cuts at NIH” have already delayed his thesis research, and these lost months cannot be recovered on a five-year stipend clock. The blow is not only logistical; it has reversed his career aspirations. Originally, Spencer was determined to serve his own government as a federal scientist. Now, he says “as an American-born U.S. citizen,” who “once aspired to work for the government as a neuroscientist” he now “desperately want(s) to leave America to pursue science in a different country.” This is a thought likely shared by many early career scientists who no longer see a future here, which is also putting America at a disadvantage for building and developing strong talent in the face of our competitors.

That sentiment is echoed by graduate student **Kenya DeBarros** from Morehouse School of Medicine: she finished her PhD degree in September 2024, with “wide-open” prospects, only to watch them evaporate “within a few months.” The choice she now faces - to shift to industry or leave the country - embodies the growing risk for U.S. science: an involuntary exodus of newly-minted PhDs at precisely the career stage when they could be entering academic pipelines, starting independent laboratories, and having the opportunity to mentor the next generation of scientists. When asked about the future of science, she stated that “it is very disheartening” that she “worked so hard for so long for the chance to make a difference in science, and a political agenda is decidedly in her way when just trying to help people is NOT a political issue.”

Additionally, an **anonymous** PhD student from University of Virginia described his selection for a position at Oak Ridge National Lab, which led to “required freeze hiring due to budget cuts and resulting shortfalls” which is hindering talent from pursuing research at the national labs due to a reduced budget. He also cites additional examples of canceled grants that would have supported his work, given his advisor “had an EPA grant which was canceled” which would have allowed him to “use one semester’s worth of funding to take the time needed to wrap up [his] research and/or give [him] some buffer to figure out next steps.” Although this is only one quote, it is likely not unique and showcases how graduate students at national labs are impacted by these funding cuts, in addition to universities, and that these actions are hindering the ability of talented students at the national laboratories to pursue research careers.

When early-career scientists at multiple levels internalize the message that their research talents and knowledge are not valued by the Trump administration when funding for their project is being cut, and that their scientific service to the greater good is contingent on political factors such as politicizing federal grantmaking, many will simply take their creativity and training elsewhere, including their contributions to the research workforce, thereby leaving the United States at a competitive disadvantage on the global stage.

### Postdoctoral Fellows: The Collapsing Bridge to Independence

If graduate students feel their careers derailed, postdoctoral fellows see the bridge ahead and their future in science collapsing, an increasingly unattainable dream. These researchers ^36^ occupy the narrow, precarious position between trainee and independent investigator. Therefore, funding instability strikes at both their livelihoods and their paths to faculty positions. In response to our survey, **Gitanjali Gnanadesikan**, a postdoc at Emory University, bluntly stated that “these drastic and arbitrary cuts to science funding hurt early career folks the worst.” She saw her NIH-funded IRACDA fellowship terminated because its DEI mandate “no longer align(ed) with NIH priorities.” The decision not only revoked her salary, it undercuts IRACDA’s mission to train diverse scientists while promoting teaching at minority-serving institutions, like Spelman College. This revocation shrinks both the biomedical workforce and the next generation of undergraduates entering it due to lack of training opportunities for talented students that would be supported by these grants. When asked about her views on the broader future of science, she stated that “the future of science looks bleak,” but that she is “trying to be hopeful that sustained advocacy and activism will be successful and improve the outlook.”

Similarly, at Arizona State University, an **anonymous** postdoc recounts the abrupt cancellation of her NSF grant in the middle of a very fulfilling role, a project “terminated early, not because it lacked value or impact, but because it no longer aligned with shifting federal priorities around equity and public-interest tech.” As to how this made her feel, she said the following “it was crushing—not just because I lost my role, but because the work mattered.” Her experience reflects two systemic dangers specifically targeting postdoctoral fellows. First, postdoctoral researchers traditionally have few safety nets, and when a single award disappears, entire research trajectories vanish with it. This is often true for under-resourced institutions like HBCUs, MSIs, and TCUs, which rely heavily on federal funding and lack large endowments or alternative funding streams. In such cases, funding cuts can threaten the institution’s very survival. Second, the projects most vulnerable to political influence are often those that challenge the status quo and/or serve marginalized communities that need this funding most. These scientific lines of inquiry are the ones the United States can least afford to lose. Samantha stated that, “we need better support for early-career researchers and more stable investment in science that serves the public—especially when it challenges the status quo” which is likely a widespread opinion among early career researchers today.

Across pre- and post-doctoral training, testimonies from survey respondents revealed a population-wide pattern: trainees are not just enduring a temporary budget squeeze. They are also evaluating whether the U.S. still affords them a viable, stable, and values-aligned space in which to build and grow their scientific careers. Without targeted interventions like bridge-funding and visa-retention incentives, our country risks hollowing out the very workforce it relies upon for future discovery and global competitiveness.

## Advocacy Actions Taken (Survey #1)

We captured early career quotes and stories, including advocacy actions taken by survey respondents in response to the Trump administration’s actions on science (survey #1). These include: sharing experiences of research funding cuts with their peers, mentors, and professional networks; learning how to advocate; contacting their representatives; educating the general public; and attending protests and organizing related events. We then collated these advocacy actions into general categories (**Table 2**).

**Table 2.** Advocacy Actions Taken by Survey Respondents. Respondents from survey #1 self-reported advocacy actions in response to the Trump administration’s actions, which we then synthesized into broad themes for easier analysis of the categories represented.

| **Advocacy Action Quotes** | **Category** |
| --- | --- |
| After the termination of my NSF funded project and job, I have been sharing my experience with peers, mentors, and professional networks to raise awareness about how these funding cuts impact (early-career) researchers and disrupt ethically grounded, community-engaged work. | Communication |
| While I haven't engaged in formal policy advocacy yet, I believe in speaking openly about these impacts and continue to look for opportunities to make our collective voices heard. | Communication |
| My advocacy actions have been learning about the scope of the issue from other scientists, thinking of how to explain it to the general public, and using my voice to educate the people around me. | Learning to advocate, communication |
| I have attended multiple protests. I am also hoping to both conduct and help organize events through Stand Up For Science. | Protests, events |
| I have called and written [to] my representatives and urged friends and family to do the same. | Contacting representatives |

## University Resources and Congressional Support

When asked how their institution has addressed professional concerns related to the research funding cuts (Q12), respondents stated that their university provided resources to keep them informed on salary and benefits (40%), counseling services and seminars (40%), no resources (30%), with other (23.3%), provided career development sessions (20%), and access to experienced mentors and peers (16.7%) (**Figure 12**). These are important aspects given that many trainees work and train in universities, and these types of supports would be critical to enabling early career researchers to thrive in science careers.

**Figure 12.**
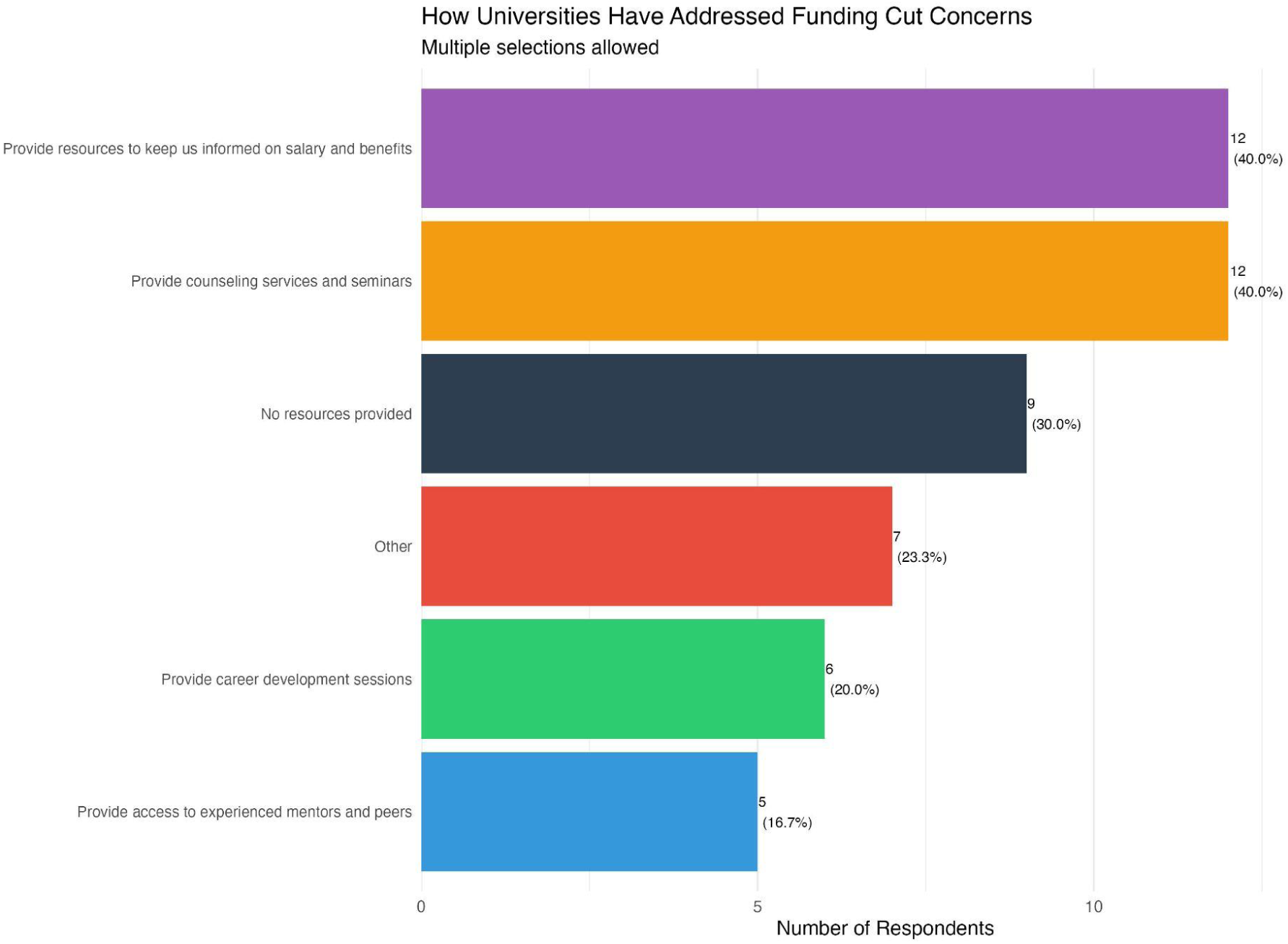
University Addressing Impacts of Federal Research Funding Cuts on Professional Concerns of Respondents (Q12). Respondents from survey #2 stated their university provided resources to keep them informed on salary and benefits (40%), counseling services and seminars (40%), no resources (30%), with other (23.3%), provided career development sessions (20%), and access to experienced mentors and peers (16.7%).

Conversely, when asked what Congress can do to help in an open-ended question (Q13), respondents largely cited the need to fund and support various systemic initiatives (including diversity, climate, health, environment) sustainably, as well as to insist that these funds are spent as intended with grants being evaluated based on scientific--not political--review, with different funding mechanisms being proposed (**Table 3**). These statements signify some confidence in the legislative branch and the fact that many Members of Congress may be supportive of early career researchers and their policy priorities at the federal level including in universities.

**Table 3.**
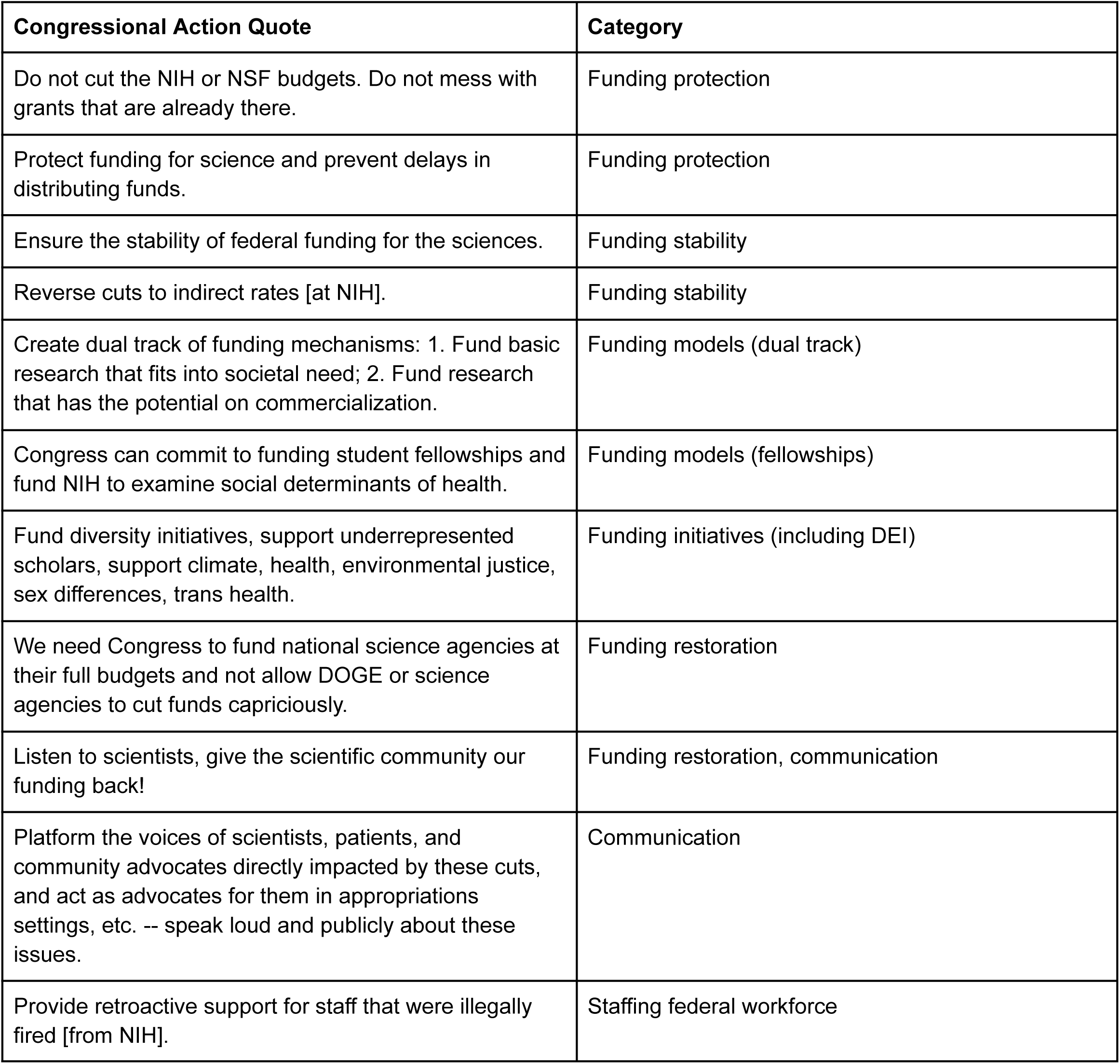
What Congress Could Do to Support Early Career Researchers (Q13). Respondents focused on funding and communication. These quotes are gathered from the data-driven survey (survey #2) with self-reported answers, with recommendations which we then categorized in broad terms due to the important role that Congress could play for the early career research pipeline. Many responses focused on NIH and NSF funding.

## Detailed Perspectives

Survey results clearly showed the detrimental impacts of the Trump administration’s actions on early career researcher respondents from various career stages, both in terms of data collection and personal narratives elaborated upon by their answers. In this section, we place the responses into a broader context and suggest potential interpretations of the results into the current and future state of the research enterprise and workforce.

The vast majority of basic scientific research occurs at academic institutions, and the lion’s share of research funding has historically come from the U.S. federal government ^37^. Our analysis captured an important segment of the STEM workforce in the early career population grappling with decreased federal research funding and impacts of these cuts on their research and careers, and beyond. Examples include removing the ability of second year graduate students to apply to the NSF GFRP ^11^ and the fact that funding for TRIO programs was set to be completely eliminated in President Trump’s FY 2026 budget proposal ^38^, as a few examples. These measures discourage talented trainees from contributing to the scientific enterprise and our global competitiveness.

This has resulted in varied impacts on universities across the nation, including likely those from our respondents (R1, R2, public, private etc.), although we did not explore this question in depth. However we speculate that, although early career researchers from certain schools or groups may have felt apprehensive to respond, many may have wanted to either express their views openly in an unbiased manner, or help the enterprise more broadly through their responses. Our analysis captured an important segment of respondents in life sciences, a large fragment of the STEM research workforce in addition to other disciplines ^39^.

Many early career respondents noted significant impacts of the funding science cuts on their professional trajectories and university experiences, which points towards the need to address their more immediate inability to complete research projects amidst uncertainty in grant funding renewals, in addition to broader issues all the way to potentially abandoning the field of research altogether. Given the impacts on more systemic issues in academia, our results show that lack of support for early career scientists from their universities can significantly derail their professional trajectories and desire to remain in the U.S. for these studies and work.

Demographic information collected from respondents, resulting in a large percentage of White individuals is also potentially notable as many academics from certain backgrounds may be apprehensive from addressing such questions ^40^ potentially due to fear of being targeted by the Trump administration. Alternatively, many have shared quotes, stories and advocacy actions taken, as an additional outlet for expression for those that may have wanted to speak out to varying degrees on either general impacts and/or also suggest advocacy actions they have taken.

Based on our analysis, universities have taken some measures to address these issues, and perhaps the responses obtained could be useful as the ecosystem seeks to rebuild from this by providing additional and/or different types of resources for early career scientists. In addition, Congress plays an important role in supporting the next generation of scientists, and opinions expressed by our respondents could suggest potential actions to be taken by the legislative branch in various facets. This could include ways to address broad systemic issues in science that could help early career scientists feel more supported in the current environment and in the future, in addition to continuously communicating the importance of this funding.

Support for early career researchers and their careers in science is imperative to continue both in the current environment and into the future, in order for the U.S. to develop a strong U.S. workforce and ensure our global competitiveness. While a number of different players have a responsibility to act on achieving this goal, trainees themselves have taken actions for their own future in science as described in the next section.

## How Early Career Researchers are Mobilizing and Community Support

In response, scientists have been fighting back through various mechanisms ^41^, and organizations have provided assistance to early career researchers such as via the American Chemical Society graduate student success grant ^42^ and Association for Women in Science microgrants for early-career research and career development ^43^.

Notably, early career researchers came together across disciplines, institutions, from multiple locations and career stages to resist and reimagine their future in science no matter where they come from. On March 7, 2025, a then-grassroots coalition and now-501c(4) nonprofit organization called Stand Up for Science (SUFS) organized a National Day of Action, with campus walkouts and rallies across 32 officially sponsored sites and over 50 solidarity events nationwide ^44^. From Ann Arbor to Albuquerque, thousands of undergraduate students, graduate students, postdoctoral scholars, faculty, and community members gathered to demand continued public investment in science. Cardboard signs stating “Courage is Contagious”, “Science is a Public Good” and “Cuts Kill Innovation” filled the streets, a public declaration that science is not only a national asset but also a democratic right.

In the aftermath, some of the SUFS organizers founded Science for Good, a nonprofit with the mission to recenter science as a tool for social good, community building, and collective problem-solving ^29^. They launched civic engagement campaigns creating casual, accessible spaces for scientists to connect with the public and share the relevance of their work. They also partnered with local breweries on *Brewing Scientists*, where researchers can co-create beer labels communicating the science happening in their communities ^45^. Each label served as a mini-billboard, elevating the work—and identities—of the researchers behind the science.

Meanwhile, other student and early-career advocacy networks launched a parallel front. The Scientist Network for Advancing Policy (SNAP) ^46^ —a coalition of over 20 student-led science policy groups—initiated the McClintock Letters Project ^47^, collecting hundreds of local newspaper op-eds and personal testimonies from scientists across disciplines, in order to put faces to the futures of those affected by science funding cuts. SNAP also wrote an open letter, echoing similar statements from the early career scientist perspective, calling on the nation’s leaders to protect the integrity of science and restore inclusive research funding. The group has since then organized 54 Congressional district visits in 29 states and designed early career curricula for science policy and advocacy.

Elsewhere, coalitions emerged on major campuses. A student-led group called Speak Out for Science (SOS) organized hands-on advocacy workshops to equip students and postdocs with the tools to write persuasive policy testimonials and defend their future ahead of appropriations and before Congress finalized its FY26 budget ^48^. Together, these efforts helped transform early-career scientists from isolated trainees into a coordinated advocacy force. They have built durable infrastructure for science policy engagement—launching new campus-based policy groups, training hundreds of scientists in advocacy skills, establishing ongoing relationships with Congressional offices, and creating open-access curricula and campaigns that can be replicated nationwide. In doing so, they have elevated early-career voices in policy debates, increased public visibility of the human impacts of funding cuts, and laid the groundwork for a more civically engaged and politically literate STEM workforce.

## Call to Action

The fight is far from over. While the March 7 actions sparked national attention, the structural damage from the FY26 budget continues to unfold. Early-career scientists—who have the most to lose and the least institutional power—remain at the forefront of both the fallout and the resistance. This includes scientists from different backgrounds, who contribute to our scientific innovation and economy and are often most vulnerable given the administration’s actions. Yet if this moment revealed anything, it’s that scientists are not only researchers—they are organizers, storytellers, coalition-builders, and defenders of democracy, and that early career scientists have a voice that matters in the process that should be heard loud and clear by our federal decision-makers. What began as a reactive moment has become a generative one: a movement for a future where science is inclusive, accountable, and deeply rooted in the public good that we can and should all participate in for our country’s future.

## Appendix

Survey questions asked during online surveys using Google Forms between May 23 and June 6, 2025.

**Survey #1:**

**Stories survey: Collecting early career researcher quotes and stories on impacts resulting from federal science funding cuts**

Email:

First and last name:

Academic institution:

Quote or story (250 words):

Advocacy action taken (250 words):

Other comments you would like to share (optional):

**Survey #2:**

**Impacts of federal science funding cuts on early career researchers**

Q1. What is your career stage?

● Undergraduate student
● PhD student
● Master’s student
● Postdoctoral researcher
● Other

Q2. How long have you been in your current position?

● Less than 1 year
● 1-2 years
● 2-3 years
● 3-4 years
● 5+ years
● Other

Q3. Which academic institution are you currently working in? (write-in) Q4. What is your field of study?

● Life sciences
● Environmental sciences
● Engineering
● Social sciences
● Other

Q5. Which of the following best describes your racial identity (select all that apply)? (optional)

● American Indian or Alaska Native
● Asian
● Black or African American
● Native Hawaiian or Other Pacific Islander
● White
● Bi or Multi-racial
● Other

Q6. Which federal agency is currently funding your research? (multiple options can be checked)

● NIH
● NSF
● DOE
● NASA
● USDA
● NIST
● DOD
● Other

Q7. On a scale of 1 - 10, how much have the recent funding science cuts affected you professionally? 1= No effect, 5= Moderately affected, 10= Severely affected

Q8. Which of the following impacts have you experienced?

● University froze hiring and/or rescinded the offer
● My stipend and benefits were cut or delayed
● Nothing has changed for me
● Other response

Q9. What is your biggest concern on potential future funding cuts impacting your research and career in science? (multiple options can be checked)

● Inability to complete research project
● Concerns with grants not being renewed
● No longer want to pursue a research career
● Concerns about long-term research job prospects
● Considering pursuing research in another country besides the U.S.
● Other

Q10. In what way have federal funding cuts affected you professionally in terms of opportunities in the following areas? (multiple options can be checked)

● Education and training
● Belonging in STEM
● Mental health and wellness
● Pay and benefits
● Other

Q11. Are you considering leaving academic research as a whole due to the funding cuts?

● Yes
● No
● Other

Q12. Has your university addressed your professional concerns as related to the funding cuts? If yes, please select your options below.

● Provide career development sessions
● Provide access to experienced mentors and peers
● Provide counseling services and seminars
● Provide resources to keep us informed on salary and benefits
● Other
● No resources provided

Q13. What can Congress do in order to support your future in research? (write-in)

## Disclaimer

This publication was written in the authors’ personal capacity and does not represent the views of their employers or organizations that the authors are or have been previously affiliated with, including any associated legislative bodies as noted in affiliations.

